# Additive-input encoding fails operator-level target-held-out prediction on current Perturb-seq screens

**DOI:** 10.64898/2026.09.30.755776

**Authors:** Kyler Fullmer, Dalton Kutzen, Tommy W. Terooatea

## Abstract

Pooled Perturb-seq screens are often interpreted through a linear steady-state model in which each targeted gene supplies an additive perturbation input. Although this formulation identifies an operator on the observed response subspace in closed form, identifiability alone does not establish that inferred operators generalize to new perturbation targets. We evaluated fixed program-space additive inputs in three Perturb-seq screens — Replogle K562 essential, Replogle RPE1 essential, and Jost 2020 — using target-grouped nested cross-validation and matched signal-to-noise linear-truth controls. For inverse operator prediction, real held-out error remained at the predict-zero baseline (0.96, 1.00, and 1.00), whereas matched linear controls achieved 0.18, 0.53, and 0.72. This discrepancy persisted across random K562 target panels, an overdetermined Jost dimension sweep, direction-only fits, ridge and truncated-SVD estimators, and five linear ground-truth ensembles. Forward target-held-out prediction was more nuanced: fixed, footprint, and learned linear encodings modestly beat a predict-training-mean baseline in K562 but not in RPE1 or Jost, and the K562 gains recovered only about 7% of the gap to the matched linear control. On the tested screens, fitted fixed program-space operators should not be interpreted as validated target-generalizing regulatory operators. We release anchor-op, a reproducible toolkit for target-grouped evaluation, identifiability checks, and matched-geometry positive controls for perturbation-response models.

## 1. Introduction

Pooled Perturb-seq screens deliver an intervention→response map at genome scale by pairing CRISPR-based perturbations with single-cell RNA-seq [1]. A common modeling step reads these responses through a linear settled-state model of regulatory dynamics: knocking down gene *g* at efficiency *κ*`_g` shifts program-space state by Δ`z_g = −J`^−1^`u_g`, where the perturbation input is `u_g = −`*κ*_g W^T^*δ*`_g`, with `W` a control-derived program basis and *δ*`_g` a one-hot on gene *g*. Under this additive-input steady-state assumption, stacking guides gives a sensitivity matrix `S = −J`^−1^`U`, and regularized inversion returns the operator’s action on the identified response subspace, `J`⋅`P_X = −U`⋅`S`^+^ with `X = range(S)`. Because this action is closed-form, the pipeline that computes it can be run on any Perturb-seq screen with a defined program basis and a calibrated efficiency estimator.

Steady-state perturbation inference has a lineage in the systems-biology literature that predates Perturb-seq: modular response analysis and reverse-engineering methods reconstruct network structure from small numbers of steady-state perturbations under an assumed linear response [3–5]. Recent Perturb-seq–era methods extend that program to genome-scale interventional data with linear or linear-cyclic structural models in gene space, including instrumental-variable, DAG, and latent-linear approaches [6–9]. At genome scale, projecting responses onto a low-dimensional program basis is the natural reduction, and `u_g = −`*κ*_g W^T^*δ*`_g` is its direct program-space form. This paper tests whether that reduction preserves target-held-out predictability. A recent benchmark separately found that a simple linear model predicts unseen single-gene perturbations in Replogle-type screens about as well as current deep-learning models [10] — a finding that motivates the learned-encoding comparator in §2.6. Answering the primary question requires a fold-honest evaluation: fit `A = −U`⋅`S`^+^ on training guides, predict `U_test = −A`⋅S_test on held-out guides, and compare the resulting held-out *ρ* to a positive control that runs the same procedure on a linear ground truth simulated at the dataset’s own guide inputs, efficiency, and per-entry noise. Under any additive-input linear model the positive control must beat the predict-zero baseline of *ρ* = 1; otherwise the identifiability calculation is not informative on the data at hand.

We ran that control on three current Perturb-seq screens: the essential-gene libraries in Replogle et al. 2022 (K562 and RPE1 lines) [1] and the titrated dose-response library in Jost et al. 2020 [2]. On the inverse task — the operator fit `A = −U`⋅`S`^+^ and the identifiability calculation that stands behind it — the fitted encoding fails to beat the predict-zero baseline on held-out prediction across all three screens, while the matched-SNR linear-truth control does. The failure of the operator-level reading is not a noise-level artifact: at the observed signal amplitude the same estimator on a linear ground truth cleanly recovers the interaction structure of the operator. It is not created by the widely used convention of selecting the 200 targets with the most cells: across 50 random 200-target K562 panels drawn from a single measurement of 1,581 qualifying targets, the fraction with real *ρ* below 0.95 is 0 / 50 (§2.3). It is not created by identifiability under-determination on Jost’s smaller target set: a d-sweep truncating the coordinate system to a comfortably overdetermined regime leaves real held-out *ρ* at the predict-zero floor. And it is not created by mis-scaling of the perturbation-strength estimate: column-normalizing the design before the fit removes all per-guide scale that a *κ* error could carry, and held-out *ρ* under the same nested cross-validation stays at the predict-zero baseline on all three screens (0.97, 1.00, 0.99) while the matched linear truth reaches 0.43, 0.78, and 0.79.

The *forward* prediction task, calibrated by the required predict-training-mean baseline and evaluated under a preregistered comparator panel of three linear encodings, gives a more nuanced per-screen picture: on K562 essential all three tested encodings — fixed, footprint, and a training-fold learned linear encoding in the style of Ahlmann-Eltze et al. 2025 [10] — beat the training-mean baseline by ~ 2 outer-fold SDs (§2.6, Table 3); on RPE1 essential and Jost 2020 none of the three beats the baseline. The K562 forward win is real and preregistered but modest: the fitted encodings close only ~ 7 % of the training-mean-to-linear-truth gap that a well-posed linear operator would deliver at matched SNR. The narrowed operator-level failure claim in the title refers to the inverse task, where the identifiability picture is unchanged on every screen.

Two candidate explanations remain live. Under the additive-input encoding, projecting the target-direction one-hot onto d = 30 control-derived programs leaves ~99% of the target direction outside the retained subspace, so the encoded input may carry little of the response-relevant signal — a *projection* failure. Independently, the linear settled-state assumption may be the wrong shape for saturating or threshold-dominated dose responses that govern actual regulatory dynamics — a *dose* failure. We describe the tests that would distinguish them and leave both to future work.

Two subsidiary correctness issues bear on any tool in this space and are handled in the software. Eps-based numerical rank on collinear guide libraries can silently declare full rank; the identifiability gate now refuses to declare full-domain recovery unless both the response and the input have full rank. And knockdown-efficiency estimators can be applied outside their valid data format; the efficiency router chooses `mean_ratio` on count data and a `detection_rate` proxy score on pre-scaled residual data, with the second labelled as a signed distributional shift rather than a fractional-knockdown estimate. anchor-op implements the pipeline, the identifiability discipline, the efficiency-routing regime, the matched-geometry positive control, and the target-held-out nested cross-validation as a reusable set of diagnostics. All numeric claims in this manuscript are round-form values; the exact values, the JSON they were computed from, and the script that produced each are listed in a machine-readable provenance table (Code availability).

## 2. Results

Box 1 (metrics used throughout §2).

- Inverse *ρ* — held-out inverse-task error: *ρ* `=` ‖`A`⋅`S_test + U_test`‖`_F /` ‖`U_test`‖`_F`, where `A = −U`⋅`S`^+^ is fit on training targets. Lower is better; *ρ* `= 1` at `A ≡ 0`.
- Forward *ρ*_fwd — held-out forward-task error: *ρ*`_fwd =` ‖*Ŝ* Ŝ_`test − S_test`‖`_F /` ‖`S_test`‖`_F`, where *Ŝ_* `test = B`⋅`U_test + b` is fit on training targets. Lower is better; *ρ*`_fwd = 1` at *Ŝ* `≡ 0`.
- Predict-zero baseline — the constant prediction `A ≡ 0` (inverse) or *S* `≡ 0` (forward). Both give *ρ* = 1 by construction. Beating predict-zero requires only that the encoding carry any predictive signal at all.
- Predict-training-mean baseline (forward only) — the constant prediction *Ŝ* `≡ b`, with `b` the per-gene mean of training responses. Beating predict-training-mean requires the encoding to carry signal *beyond* the shared response mode captured by `b`, and is the calibrated forward-task criterion used from §2.6 onward.

**Figure 1.**
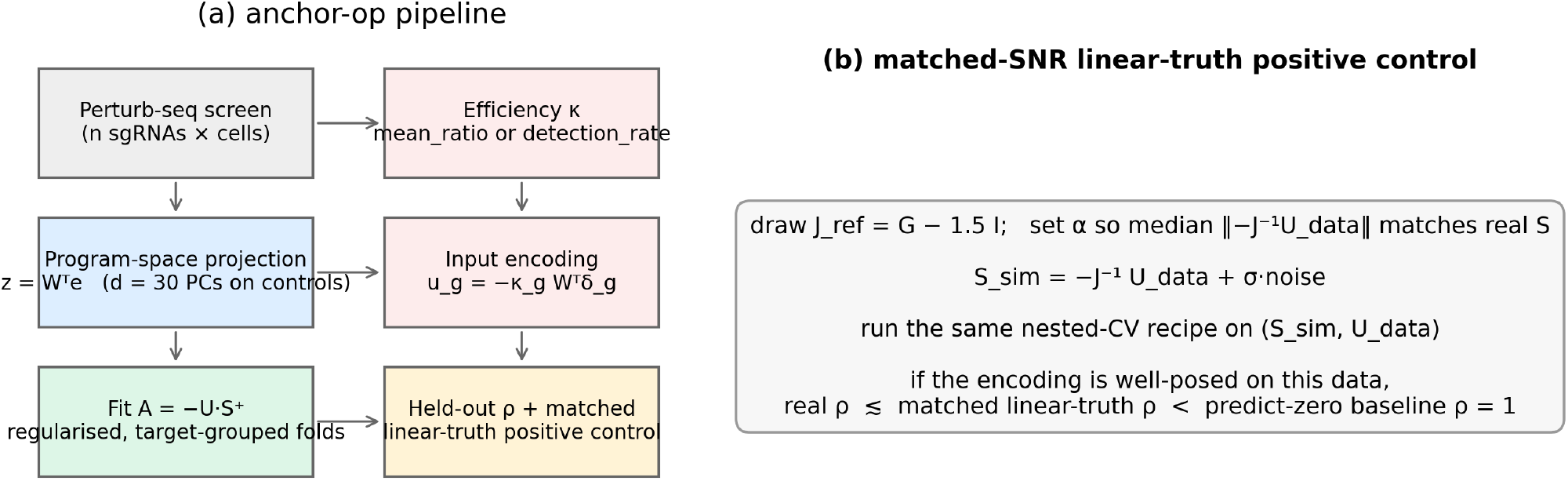
*(a)* anchor-op fitting pipeline: program-space projection of expression data on a control-derived PCA basis, encoding of each perturbation input, and a regularised target-grouped-fold operator fit. *(b)* Matched-SNR linear-truth positive control used throughout: simulate a known linear ground truth at the dataset’s own U, *κ*, and σ, then run the same nested-CV recipe on the simulated (S, U) pair. If the encoding is well-posed on this data, real *ρ* should track the matched-linear-truth *ρ* below the predict-zero baseline of 1.

### 2.1 Target-held-out prediction fails against the matched linear control

For each screen we fit the additive-input operator `A = −U`⋅`S`^+^ using truncated SVD, with the retained rank chosen by nested cross-validation inside each outer training fold (Methods). Both outer and inner folds are target-grouped, so no sibling sgRNA leaks between train and test. The rank grid includes zero, which represents the predict-zero baseline `A ≡ 0` and gives held-out *ρ* = 1 exactly; nested CV therefore selects “shrink to zero” whenever the fitted model does worse than predict-zero on inner validation.

We report pooled held-out *ρ* with its across-outer-fold standard deviation on the real data, and mean ± SD across 15 independent matched-SNR linear-truth simulations. Table 1 summarizes. Table 1. Target-held-out *ρ* under nested cross-validation.

**Table 1.**
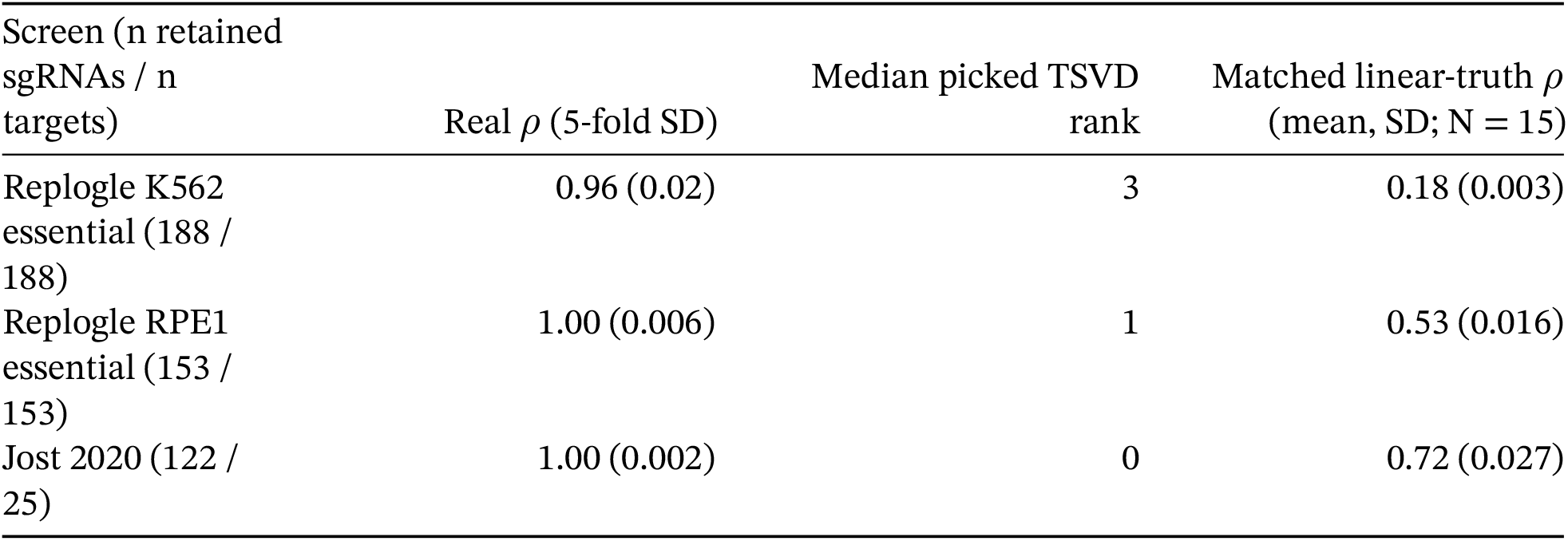
Target-held-out *ρ* under nested cross-validation.

**Table 2.** Jost d-sweep under target-grouped nested cross-validation.

| d | Real $\rho$ (pooled) | Median picked rank | Matched linear-truth $\rho$ (mean, SD; N = 15) | Regime |
| --- | --- | --- | --- | --- |
| 5 | 1.01 | 0 | 0.32 (0.038) | overdetermined |
| 10 | 1.00 | 0 | 0.85 (0.067) | overdetermined |
| 15 | 0.99 | 2 | 0.46 (0.031) | overdetermined |

| d | Real $\rho$ (pooled) | Median picked rank | Matched linear-truth $\rho$<br>(mean, SD; N = 15) | Regime |
| --- | --- | --- | --- | --- |
| 20 | 1.00 | 0 | 0.55 (0.061) | overdetermined |
| 25 | 1.00 | 1 | 0.69 (0.037) | underdetermined |
| 30 | 1.00 | 0 | 0.73 (0.043) | underdetermined |

**Figure 2.**
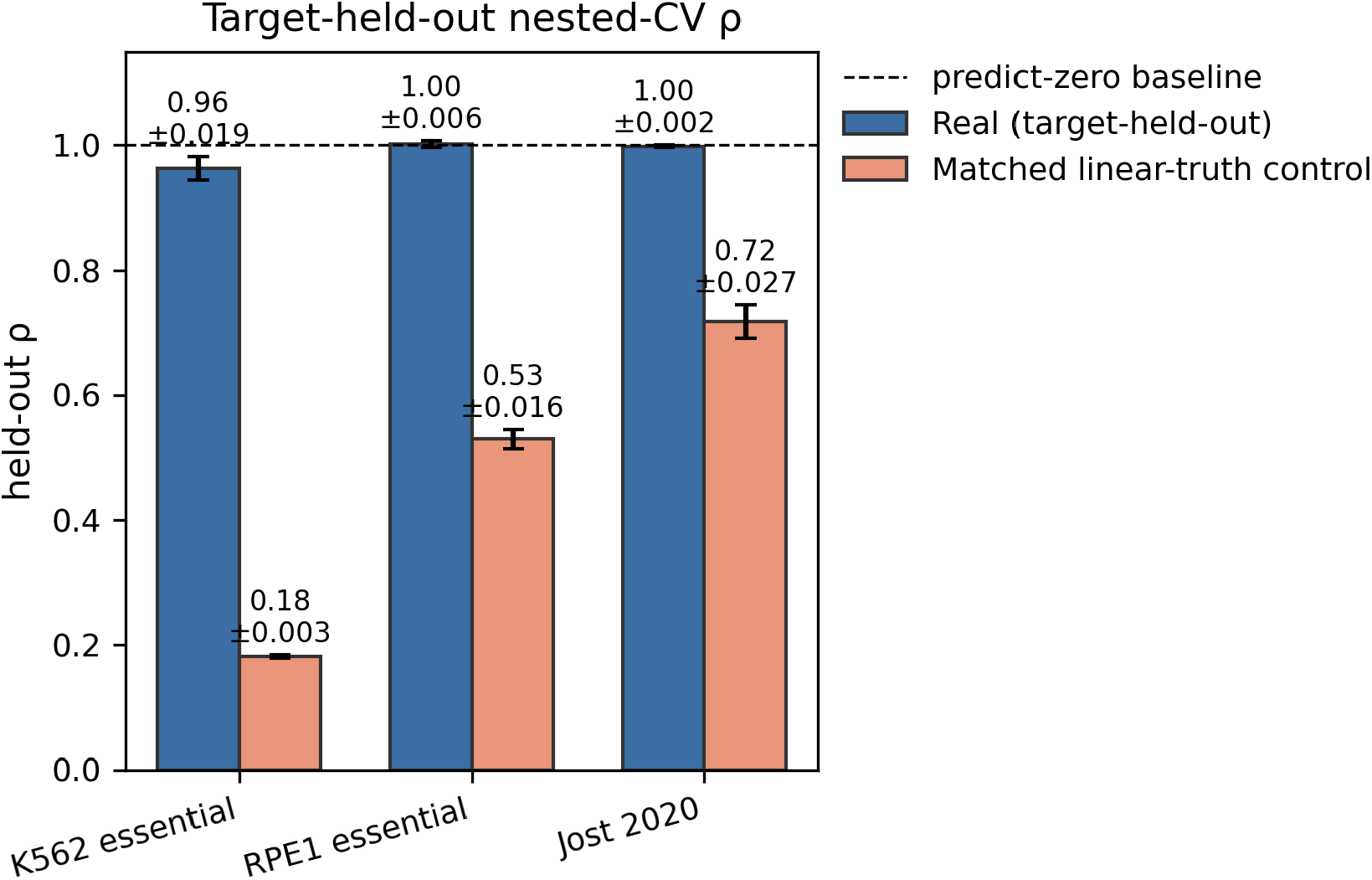
Target-held-out nested-CV held-out *ρ* on the three screens (real, blue; matched linear-truth control, orange). Error bars: 5-fold SD for real, cross-replicate SD across 15 sims for the control. Dashed line: predict-zero baseline *ρ* = 1.

Every real *ρ* sits at or above the predict-zero baseline within one fold-level SD. On K562 the median picked rank is 3; on RPE1 it is 1; on Jost it is 0 — nested CV picks “shrink to zero” as the best per-fold model, so the fitted operator loses to no model at all. Under the same estimator on the same U and per-entry σ, a matched-SNR linear ground truth reaches held-out *ρ* well below the predict-zero baseline in every case.

The three screens differ in library size and design. Replogle-essential libraries retain one sgRNA per target after the released target-level aggregation step. Jost carries several sgRNAs per target at graded activities, spanning *κ* ∈ `[0.07, 1.00]` with count-based estimates from `mean_ratio` on UMIs — roughly twice Replogle’s *κ* range in a design purpose-built for dose response — and 25 targets total. Their per-entry noise anchors, estimated from within-guide cell-level split-half bootstraps in each screen’s own basis, are 0.24 (K562), 0.35 (RPE1), and 0.066 (Jost, per sgRNA). The failure is not restricted to a specific library size, design, or noise level.

### 2.2 The failure is not a noise problem

At the amplitude of the released simulation ensemble the noise-free response is much smaller than the observed response — a factor of ~360 on K562 by median column norm. At that amplitude the signal-to-noise ratio is low and any TSVD estimator, linear-truth or not, has almost no chance of recovering the operator. A signal-scale sweep varying the multiplier α`_S` on the noise-free response confirms this and identifies the α_S that reproduces the observed median column norm of `S`.

At the data-matched amplitude the same TSVD estimator on a linear ground truth cleanly recovers the operator: full-matrix Frobenius cosine 0.99 (K562) and 0.91 (RPE1) at N = 200 replicates. Those full-matrix cosines are dominated by the shared diagonal shift carried by every replicate in the ensemble; their cross-replicate null is around 0.70. The *interaction-only* cosine — computed after subtracting the diagonal from both `A` and `J` — isolates the recoverable structure. It reaches 0.98 (SD 0.003) on K562 and 0.74 (SD 0.023) on RPE1 at matched amplitude, against a cross-replicate null of 0.003 in both cases (N = 200).

The estimator can recover a known operator’s interactions at the signal amplitude of the observed response. The gap between “an estimator that recovers a linear operator’s interactions at matched SNR” and “an estimator that fails held-out prediction on real data at the same SNR” is the point. Under the additive-input encoding, the model fits `A` and predicts `U = −A`⋅`S` on held-out guides. On the linear-truth positive control that prediction succeeds; on the real data it does not. The failure lies somewhere between the data-generating process and the additive-input model, not in the fitting machinery.

The finding is not specific to the dense ground-truth ensemble either. Rerunning the matched-SNR linear-truth control at each screen’s own amplitude across five ensembles — dense, sparse-10%, sparse-2%, rank-5, and block-modular (Methods; Supplementary Fig S2) — leaves the “linear truth beats predict-zero at matched SNR” conclusion intact on every (screen × ensemble) cell (15/15). No ensemble tested flips the sign of the conclusion.

**Figure 3.**
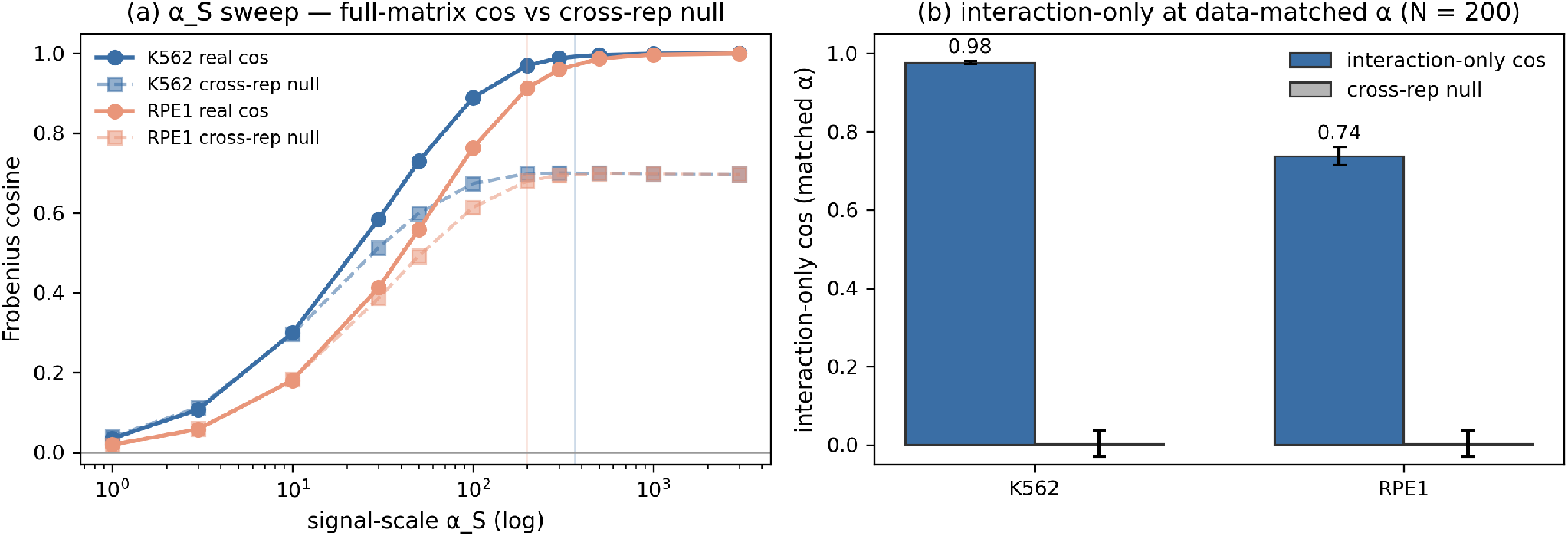
*(a)* Signal-scale α_S sweep of full-matrix Frobenius cosine on K562 and RPE1, with the cross-replicate null on each curve. Vertical bars mark the data-implied 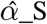. *(b)* Interaction-only cosine at data-matched amplitude (N = 200) against the cross-replicate null on the same replicates. The estimator recovers a known operator’s interactions cleanly at the observed signal amplitude.

### 2.3 The failure is not target selection, input dimension, or *κ* mis-scaling

Selection bias. The publicly released essential-gene notebooks pick the 200 targets with the most cells after a minimum-cell filter. In essential-gene screens, high cell count correlates with mild fitness defect, so the selection preferentially retains weakly-affected targets. To rule out the possibility that the failure is specific to one favourable target set, we measured all K562 targets that pass the pipeline’s retention filter on the full essential library in one fit (1,581 targets; Σ`_ctrl` and program basis saved) and then drew 50 random 200-target panels from that single measurement. Preregistered summary: the fraction of panels with real nested-CV *ρ* below 0.95. The distribution is tight and above 0.95 in every draw — real *ρ* median 0.9988, 5th–95th percentile 0.9855–1.0054, minimum 0.9698 across 50 panels — while the matched linear-truth *ρ* under the same recipe (per-panel mean over N = 5 sims) has median 0.3700 and 5th–95th percentile 0.3136–0.4416. The preregistered fraction is 0 / 50 (0%). The failure is a property of the target-held-out prediction task on this screen, not of any particular random 200-target draw.

Design underdetermination. Jost has 25 target directions and d = 30 program dimensions; under 5-fold target-grouped CV each outer training set spans only ~20 target directions, which is fewer than d. If the encoding failed on Jost only because of this under-determination, the failure should disappear when the coordinate system is truncated to a smaller d where training targets comfortably exceed d. We swept d ∈ {5, 10, 15, 20, 25, 30} on Jost, recomputing α_S at each d so the matched linear-truth control tracks the same signal amplitude at each point.

At d = 5, where the identifiability regime is comfortably overdetermined, the matched linear-truth *ρ* is 0.32 while real Jost *ρ* is 1.01. The matched linear-truth *ρ* varies non-monotonically with d — from 0.32 at d = 5 to 0.85 at d = 10 — because α_S is recomputed at each d and different truncations of the coordinate system carry different fractions of the observed response amplitude; but it stays below the real *ρ* at every d, so the encoding fails independent of the identifiability regime.

*κ* mis-scaling. The Replogle-essential pipeline uses a residual-space `detection_rate` proxy for *κ* (Methods), which is a signed distributional-shift statistic rather than a fractional-knockdown estimate. If mis-scaled per-guide *κ* were the source of the failure, column-normalizing S and U before the fit — which removes all per-guide scale from the fitting problem — should improve held-out *ρ*. It does not. Under the same target-grouped nested cross-validation, direction-only real *ρ* is 0.97 (5-fold SD 0.007) on K562, 1.00 (SD 0.002) on RPE1, and 0.99 (SD 0.017) on Jost. The matched linear-truth control on the same column-normalized design reaches 0.43 (SD 0.018) on K562, 0.78 (SD 0.013) on RPE1, and 0.79 (SD 0.030) on Jost, all well below the predict-zero baseline. Column-normalizing raises the matched-linear *ρ* on RPE1 and Jost — the transform discards real per-guide scale information those screens’ *κ* estimators carry (Replogle-essential’s residual-space proxy is uninformative in this regard; Jost’s count-based `mean_ratio` is informative) — but *κ* mis-scaling cannot on its own account for the encoding’s failure on any of the three screens.

**Figure 4.**
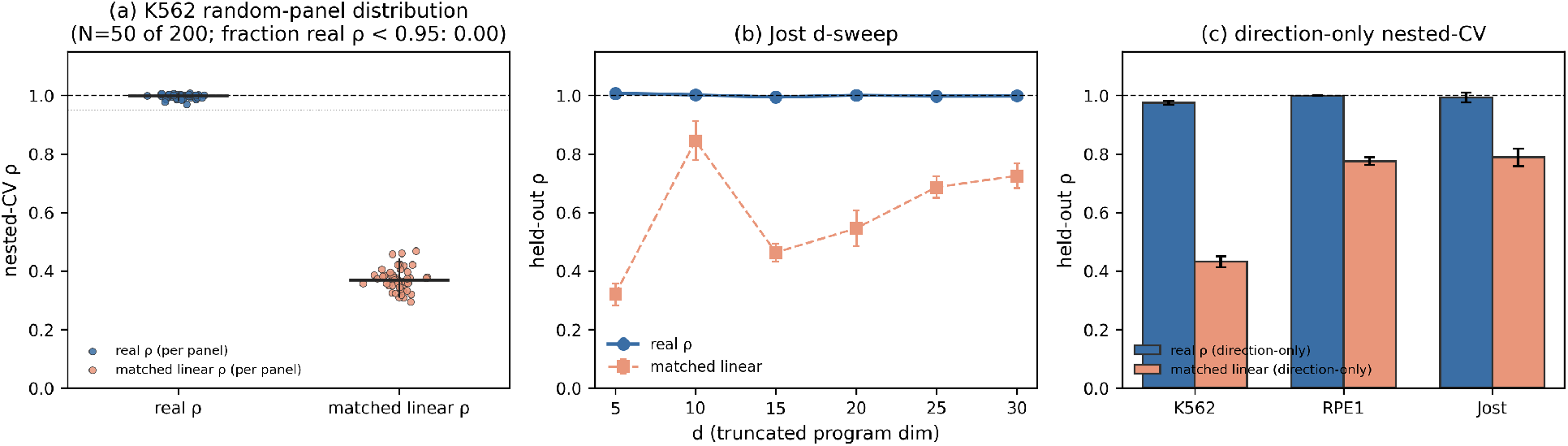
Robustness checks. *(a)* K562 random-panel distribution: 50 random 200-target draws from a single measurement of the 1,581 qualifying targets. Real nested-CV *ρ* (blue) has median 0.9988 with 5th–95th percentile band 0.9855–1.0054 (fraction below 0.95: 0 / 50). Matched linear-truth *ρ* (orange, per-panel mean of N = 5 sims) has median 0.3700 with band 0.3136–0.4416. *(b)* Jost d-sweep under target-grouped nested CV: real *ρ* stays at the predict-zero baseline across d ∈ {5, 10, 15, 20, 25, 30}, while the matched control drops well below 1 in every regime. *(c)* Direction-only refit under the same nested-CV recipe on all three screens rules out *κ* mis-scaling.

### 2.4 The pipeline captures within-target dose interpolation on Jost

Jost 2020 pairs each of its 25 targets with 3–6 sgRNAs at graded activities. Under target-held-out prediction (the setting in §2.1–2.3) a held-out target’s sgRNAs have no siblings in training, and the model is asked to predict a new perturbation direction. Under *guide-level* folds — where every held-out sgRNA has siblings for its own target in training — the model is instead asked to interpolate amplitude along a *known* direction. It predicts held-out `U_test` given `S_test` along that known direction. That interpolation *ρ* is 0.66 on real Jost, against 0.22 (SD 0.008, N = 15) for the matched linear-truth control on the same guide-level folds (Supplementary Fig S1, left).

Two things follow. The pipeline does capture within-target amplitude structure when a target’s direction is available: 0.66 is meaningfully below the predict-zero baseline of 1 and well above the null. But the amplitude the pipeline captures is not operator recovery. Under any additive-input linear model, every guide for one target moves the system along the same direction — *κ*`_g` rescales columns, W^T^*δ*`_g` is fixed — and the guide-level fold split reduces the prediction task to one that a per-target one-parameter fit could pass. We label the guide-level Jost result “within-target dose interpolation along a known target direction” and keep it separate from the target-held-out claim that governs the main result.

### 2.5 A widely used linearity threshold is uncalibrated at these signal-to-noise ratios

The `rel_diff` statistic splits guides at the median efficiency, fits an operator per half, and returns the Frobenius-normalized difference on the shared identified subspace. A prespecified threshold `rel_diff` ≤ `0.25` was chosen at preregistration to declare the additive-input model consistent with the data. Real data on Replogle gives `rel_diff` = 1.47 (K562) and 1.57 (RPE1); on Jost it is 1.26. Every screen fails the threshold. But so does a matched linear ground truth at each screen’s own signal amplitude and noise: 0.33 (SD 0.01) on K562, 0.98 (SD 0.02) on RPE1, and 1.05 (SD 0.03) on Jost (N = 15). On RPE1 and Jost even a perfectly linear generator fails the 0.25 threshold; only on K562 does it drop below 1.0. It is miscalibrated (Supplementary Fig S1, right). The calibrated comparison — real vs matched linear truth at each screen’s own SNR — corroborates the target-held-out failure on all three: real `rel_diff` exceeds matched linear-truth `rel_diff` by ~1.14 (K562), ~0.59 (RPE1), and ~0.21 (Jost, about 7 linear-truth SDs above the matched-control mean).

We ship a separate correctness fix at the identifiability level: the gate that labels a measurement “full-domain identified” now requires both the input rank and the retained response rank to equal d at the preregistered tolerance. Jost’s input rank is 24 at that tolerance, so the gate now correctly labels the Jost fit as partially identified regardless of what noise does to the retained response rank.

### 2.6 A comparator panel: three linear encodings succeed on K562 essential; all three fail on RPE1 and Jost

Sections §2.1–§2.5 test the fixed program-space encoding `u_g = −`*κ*_g W^T^*δ*`_g` under the inverse task (fit `A = −U`⋅`S`^+^, predict *Û*`_test = −A`⋅`S_test`). To place that failure in the context of the two natural alternatives — a fixed *footprint* encoding and a *learned* linear encoding in the style of the current Perturb-seq-prediction benchmark [10] — a preregistered comparator panel runs all three encodings under a single evaluation on each of the three screens (Methods §4.5b, §4.8), and adds a predict-training-mean baseline. About half of each screen’s response energy is a shared program-space mode, so beating predict-zero is trivial; the calibrated question is whether an encoding beats the training-mean baseline. The two-fold-SD criterion is a preregistered practical decision rule for this benchmark, not a formal hypothesis-test threshold.

The forward task is *ρ*`_fwd =` ‖Ŝ_`test − S_test`‖`_F /` ‖`S_test`‖`_F` under target-grouped nested-CV. Baselines: predict-zero (*ρ*_fwd = 1) and predict-training-mean (column mean of training `S` broadcast across the held-out targets). Encodings, each fit as *Ŝ* `= B`⋅`U + b` with `b` the training-fold mean response and `B` a ridge fit on the centered training responses (λ by inner CV; the log grid includes λ `= 1`×10^30^, which collapses `B` → `0` and yields *Ŝ* `= b`): (i) *fixed* — U_g = W^T^*δ*`_g`; (ii) *footprint* — U_g = W^T^ Σ`_ctrl` *δ*`_g`; (iii) *learned* — a learned linear embedding following Ahlmann-Eltze et al. [10] in gene space, with the gene embedding derived from a PCA of the training targets’ gene-space pseudobulk responses only, refit within every outer fold. The held-out target’s embedding is that gene’s row in the training-derived PCA — legitimate because it is a feature-gene, not a response. Predictions are made in gene space with the same training-fold-mean intercept and then projected through `W` for the program-space metric (the gene-space metric is also reported). Nulls for the learned encoding: a shuffled-embedding null (≥ 100 permutations of the target→embedding map, refit per permutation, with the intercept preserved). The preregistered success criterion is *ρ*_fwd more than 2 outer-fold SDs below the training-mean baseline AND (learned only) below the 2.5th percentile of the shuffled-embedding null.

The training-fold intercept is what makes the training-mean baseline reachable within the encoding class: under λ → ∞, `B` → `0`, so *Ŝ* `= b = training-mean baseline` exactly. A sanity call at λ `= 1`×10^30^ returns *ρ*_fwd equal to the training-mean baseline on each screen with `|gap| < 10`^−8^ (Methods §4.5b). Under this convention every encoding must beat the intercept-only prediction on inner validation to be picked by nested CV.

Table 3. Forward comparator panel (target-grouped nested-CV *ρ*_fwd; lower is better; *ρ*_fwd = 1 is predict-zero). The `2` × `SD_tm` decision threshold is 0.0436 on K562, 0.0586 on RPE1, and 0.1364 on Jost; encoding-vs-baseline gaps that exceed this threshold are marked PASS (K562 fixed 0.0475, K562 footprint 0.0485, K562 learned 0.0468), the rest are FAIL. The table is split in two for typesetting; both halves list the same three screens.

**Table 3a.**
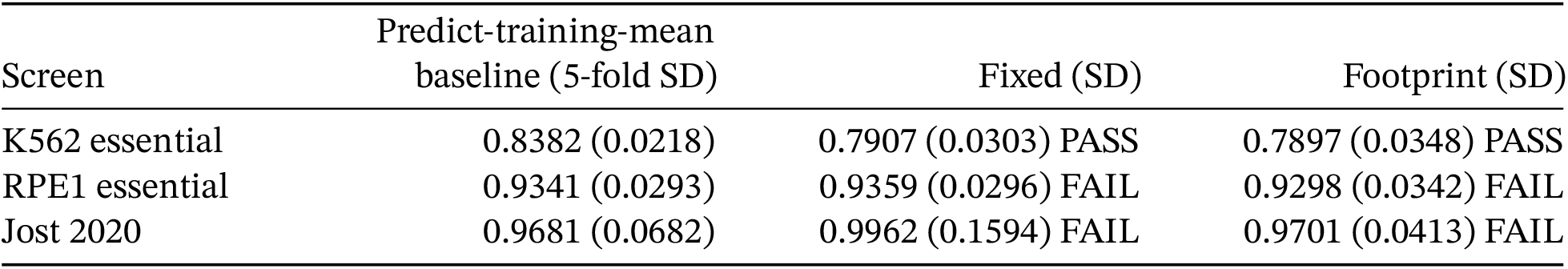
Baselines and fixed / footprint encodings.

**Table 3b.** Learned encoding (program and gene space) and its shuffled-embedding null.

| Screen | Learned prog (SD) | Learned gene | Learned shuffled-null<br>2.5 % |
| --- | --- | --- | --- |
| K562 essential | 0.7914 (0.0401) PASS | 0.8563 | 0.8366 |
| RPE1 essential | 0.9267 (0.0086) FAIL | 0.9660 | 0.9273 |
| Jost 2020 | 0.9528 (0.1024) FAIL | 0.9559 | 0.9487 |

**Table 4.**
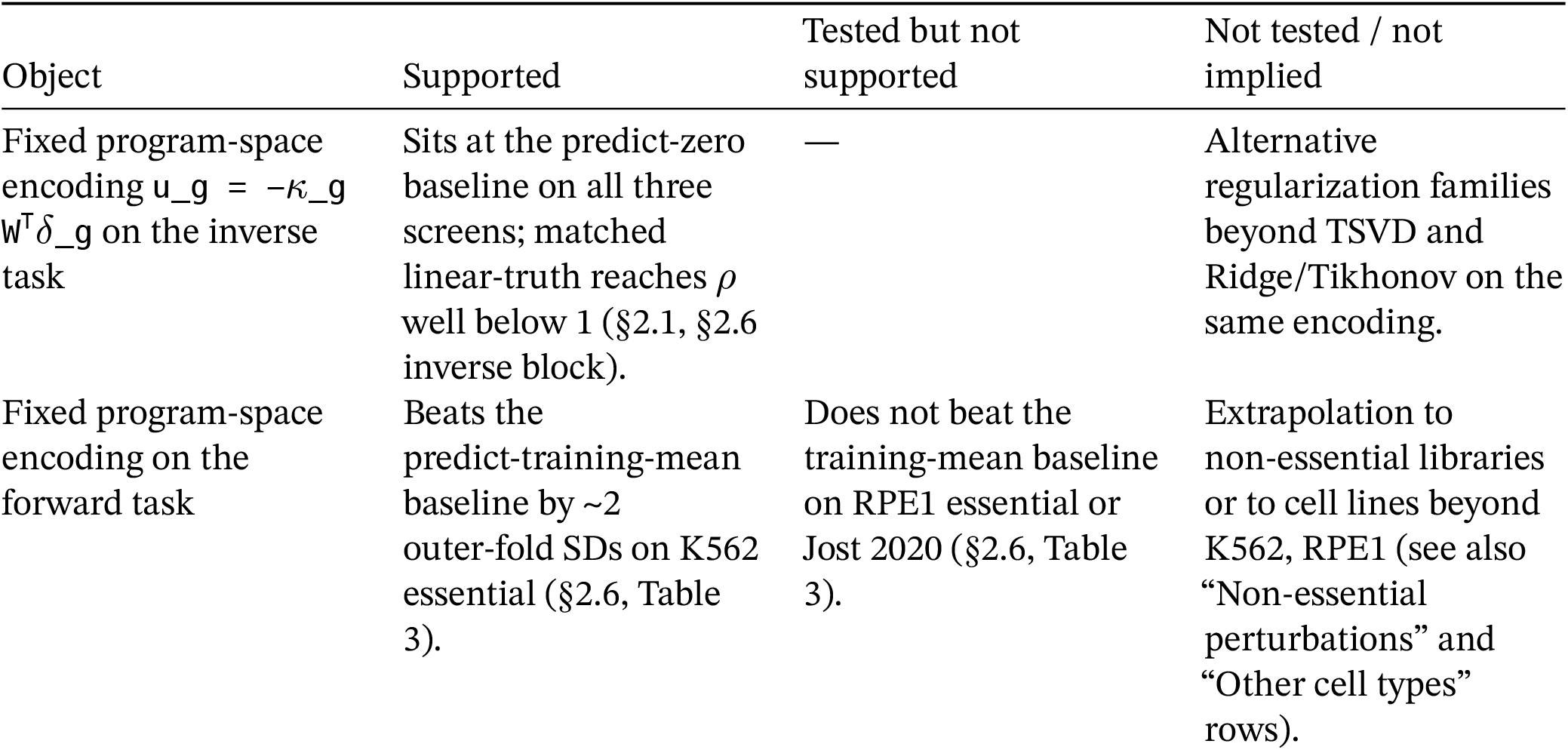
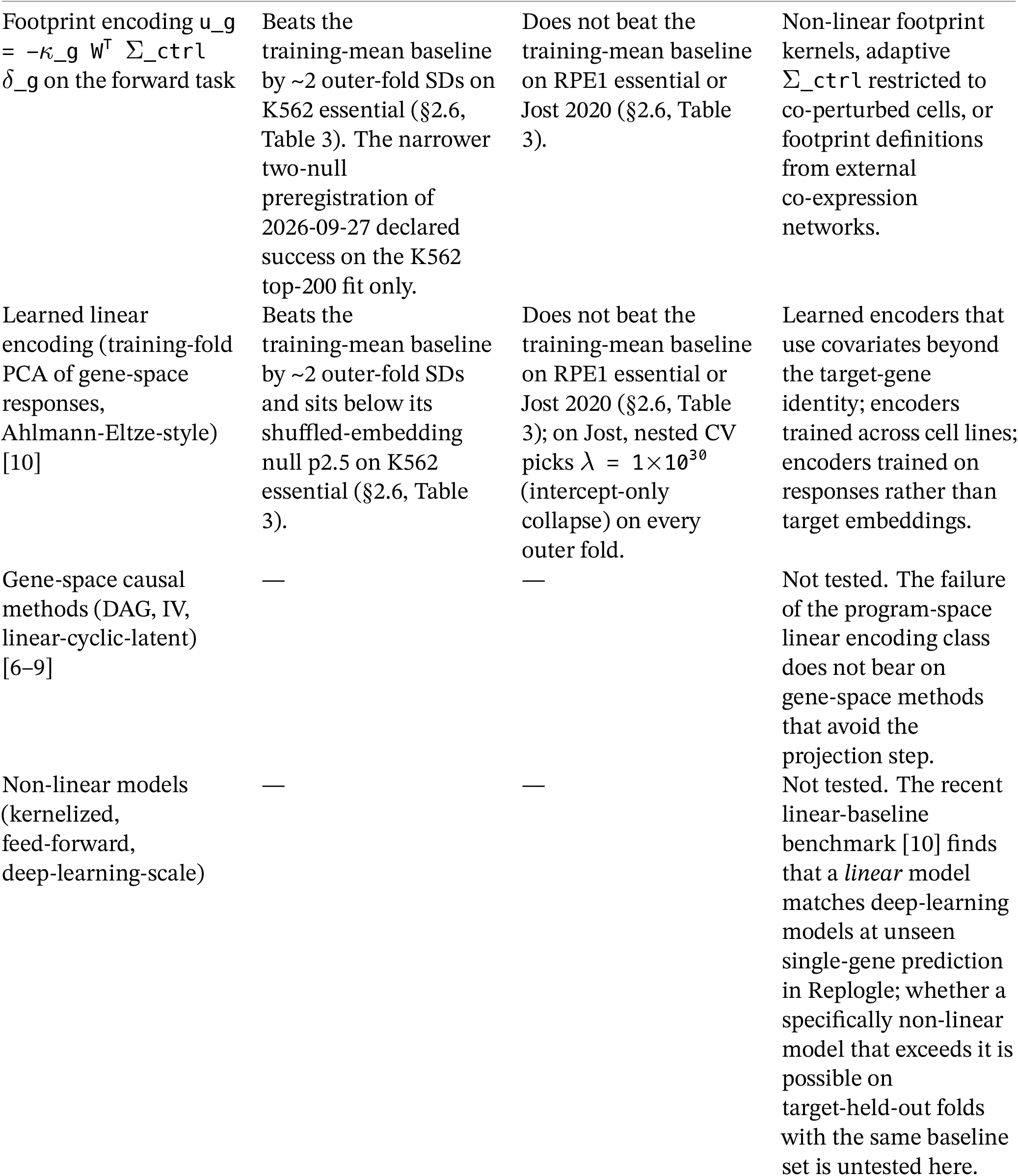
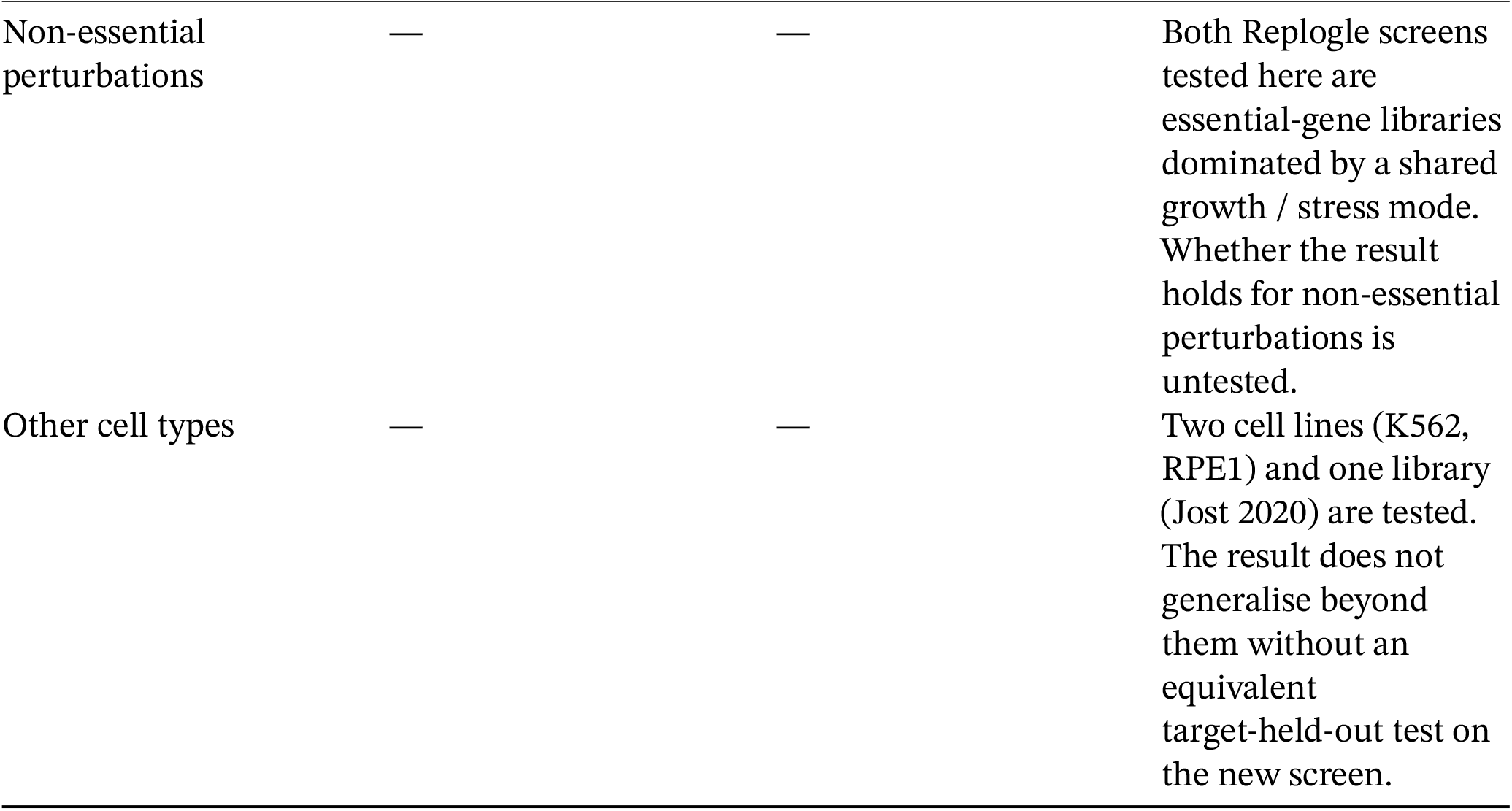
What this paper does and does not claim.

Verdict. On K562 essential all three linear encodings — fixed, footprint, and learned — beat the training-mean baseline by ~2 outer-fold SDs (gaps 0.0475, 0.0485, 0.0468 against a 2 × baseline-SD threshold of 0.0436) and the learned encoding sits below its shuffled-embedding null p2.5 (0.7914 < 0.8366). On RPE1 essential and Jost 2020, no encoding beats the training-mean baseline: the *ρ*_fwd gaps against `2` × `SD_tm` are all inside the fold-level noise. The learned encoding on RPE1 sits marginally below its shuffled-null p2.5 (0.9267 < 0.9273 by 0.0006) but does not meet the 2-SD gap requirement; the learned encoding on Jost sits *above* its shuffled-null p2.5 (0.9528 > 0.9487), and nested CV picks λ `= 1`×10^30^ (the intercept-only collapse) on every outer fold — the fitted map does not beat the training-mean intercept on any inner validation split on Jost.

Decision rule invoked. Learned succeeds on 1/3 screens (K562 only), so branch (i) of the preregistered rule (promote a learned-encoding-class claim on ≥ 2/3 screens) does not apply. Fixed and footprint succeed on 1/3 (K562) and fail on 2/3, and learned fails on 2/3, so branch (ii) (“neither fixed nor learned linear encodings predict held-out targets beyond the training mean on these screens”) does not describe the K562 outcome. The applicable branch is (iii) *mixed*: per-screen reporting, no cross-screen ranking. Table 3 is the per-screen report. No promotion of a title-level claim. The paper reports the K562 forward-task success and the RPE1 / Jost failures side by side; the operator-level (inverse-direction) reading in §2.1–§2.5 remains a failure on all three screens.

The matched linear truth under each encoding’s `U` — same recipe, linear ground truth simulated at each screen’s own signal amplitude and noise, intercept applied — beats the training-mean baseline by more than 2 SDs on every (encoding × screen) cell (K562 fixed 0.19 SD 0.03; K562 footprint 0.18 SD 0.04; RPE1 fixed 0.39 SD 0.04; RPE1 footprint 0.29 SD 0.02; Jost fixed 0.69 SD 0.04; Jost footprint 0.53 SD 0.08). The estimator recovers signal that only two of the six real (encoding × screen) cells recover. Even on K562, where fixed and footprint succeed against the training-mean baseline, the real data close only a small fraction of the training-mean-to-linear-truth gap: the training-mean baseline is 0.838, the matched-linear-truth *ρ*_fwd is 0.186, and the fitted fixed *ρ*_fwd of 0.791 closes (0.838 − 0.791) / (0.838 − 0.186) ≈ 7% of that gap (footprint closes ≈ 7%). What K562’s linear-truth generator delivers at matched SNR is a 78% reduction of *ρ*_fwd from the baseline; what the fitted encodings recover is a 5–6% reduction. The K562 success is a small but preregistered win, not an approach to what a well-posed linear operator on the same design would predict.

The inverse-direction comparator (predict `U_test` from `S_test` under TSVD and Ridge/Tikhonov, matched linear-truth alongside) tells the §2.1 story unchanged: both estimators track each other closely on the real data (K562 TSVD 0.97, Ridge 0.96; RPE1 both ≈ 1.00; Jost both ≈ 1.00) while the matched-linear-truth *ρ* reaches 0.18 (K562), 0.53 (RPE1), and 0.70–0.73 (Jost). Ridge with λ → ∞ (collapsing `A` to zero) is included in the log-λ grid and picked by nested CV whenever it beats the fitted model on inner validation — this is the shrink-to-zero baseline familiar from §2.1.

**Figure 5.**
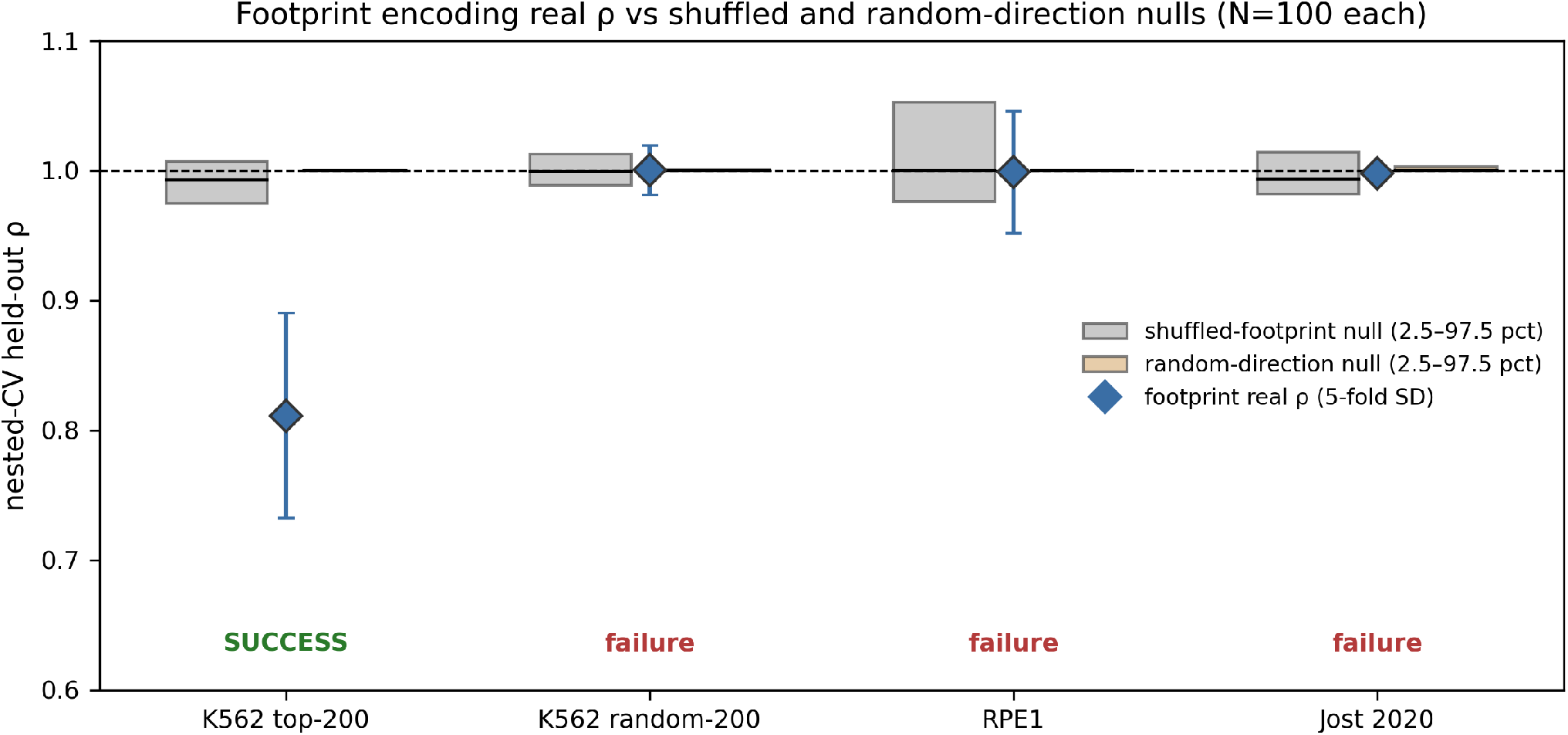
Footprint-encoding nested-CV *ρ* on the four earlier fits used in the preregistered footprint check (blue diamonds, with 5-fold SD) against the two preregistered nulls, each drawn as the 2.5–97.5 percentile band with the median as a horizontal bar: shuffled-footprint null (N = 100 permutations, grey) and random-direction null (N = 100 draws with matched column norms, tan). The K562 top-200 cell of this earlier fit is the same design in which the comparator panel of Fig 6 finds the footprint *ρ*_fwd= 0.79 on the forward task.

**Figure 6.**
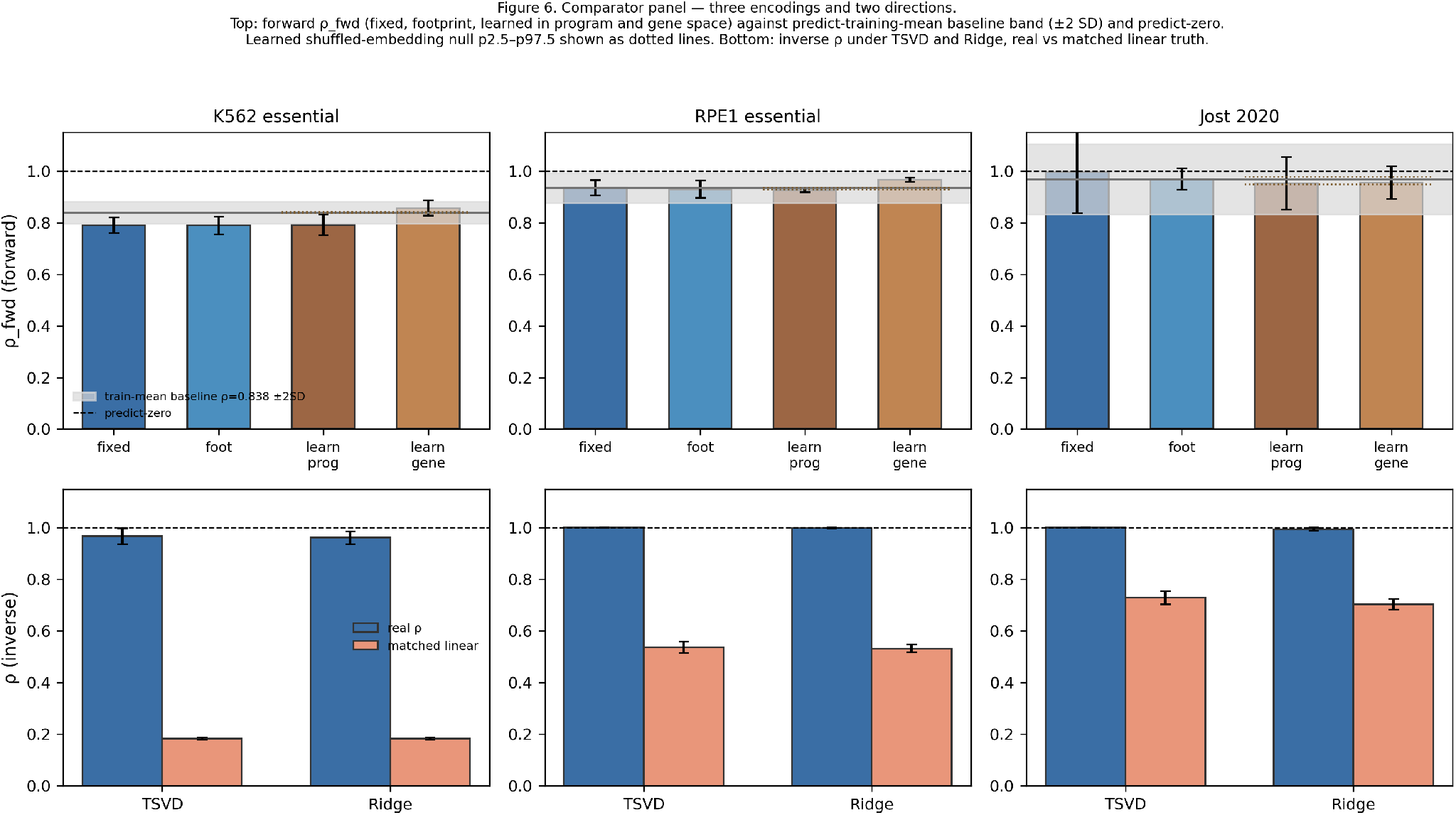
Comparator panel across the three screens. Top row: forward direction (*ρ*`_fwd` under fixed, footprint, and learned encodings, all with the training-fold mean intercept, with the predict-training-mean baseline band at ±2 × baseline-SD and the learned shuffled-embedding null p2.5–p97.5 as a dotted range). Bottom row: inverse direction (TSVD and Ridge on the real data with matched linear-truth *ρ* alongside). Dashed lines: predict-zero baseline. On K562 all three encodings sit visibly below the training-mean baseline band; on RPE1 and Jost every encoding sits inside the band.

## 3. Discussion

### 3.1 What the result says

Under three linear encodings — fixed program-space `u_g = −`*κ*_g W^T^*δ*`_g`, control-covariance footprint, and a training-fold learned linear encoding — and both directions of the operator-inference task on three current Perturb-seq screens, the target-held-out outcome is per-screen. On the inverse direction all three real-data *ρ* sit at the predict-zero baseline while a matched linear ground truth reaches *ρ* well below 1 on every screen. On the forward direction, calibrated by the predict-training-mean baseline, all three encodings beat the baseline by ~2 outer-fold SDs on K562 essential (and the learned encoding sits below its shuffled-embedding null p2.5), while none of the three beats the baseline on RPE1 essential or Jost 2020 — on Jost the learned map degenerates to the training-mean intercept on every fold selected by nested CV. The K562 forward signal beyond the training mean is modest against a matched-linear *ρ*_fwd of 0.19 — the estimator has substantially more room to move than any fitted encoding uses. Fitted spectra or edges from the fixed program-space encoding on RPE1 and Jost should not be read as quantitatively estimated operators; on K562 all three fitted maps beat the training-mean baseline on the forward task, but the inverse identifiability picture (§2.1) is unchanged and the operator-level interpretation of a fitted map remains cautious. The result does not rule out recovery under a different encoder class (nonlinear encoders that use covariates beyond the target-gene identity, or encoders trained across cell lines), a lower-dimensional target, a stronger structural prior, an assay design with true replicate targets and calibrated dose titration, or a gene-space (rather than program-space) causal method. We frame anchor-op as a calibrated evaluation and diagnostic toolkit for the additive-input linear model class on current Perturb-seq screens rather than as a new operator-inference method.

### 3.2 What is not yet decided

The comparator panel in §2.6 tests three encodings — fixed program-space, control-covariance footprint, and a training-fold learned linear encoding in the style of the current Perturb-seq-prediction benchmark [10] — under a single evaluation that includes the required predict-training-mean baseline. On K562 essential all three encodings beat the training-mean baseline by ~2 outer-fold SDs and the learned encoding sits below its shuffled-embedding null p2.5. On RPE1 essential and Jost 2020 no encoding beats the baseline; on Jost nested CV picks the intercept-only collapse (λ `= 1`×10^30^) on every outer fold, so the fitted learned map does not beat the training-mean intercept on any inner validation split. Two candidate mechanisms remain consistent with what we ran; a third live question — why K562 admits a modest predictable signal beyond the training mean and RPE1 and Jost do not — is discussed alongside.

Projection failure. The fixed additive-input encoding pushes a target-direction one-hot through a d = 30 control-derived program basis. The projected input W^T^*δ*`_g` has median column norm ~0.09 on the essential-gene screens — about 99% of the target direction lies outside the retained subspace. If the target’s response-relevant direction is not among the top d control-derived programs, the encoded input carries little response-relevant signal and the fitting task becomes ill-posed on target-held-out folds. The footprint encoding replaces the one-hot with the target’s control-cell co-expression neighbourhood; the learned encoding lets the target’s gene-space PCA row parameterise the input direction. On the forward task calibrated by the training-mean baseline, none of the three encodings rescues operator-level prediction, and their forward gains are confined to K562 and small. Whether a *different* learned encoder — a nonlinear representation, one trained across cell lines, or one that uses paired covariates beyond the target-gene identity — would rescue prediction is untested here.

Dose non-linearity. The linear settled-state assumption `S = −J`^−1^`U` may not describe the assay’s actual dynamics if the response saturates or thresholds in dose. Under any additive-linear model, all sgRNAs for one target move the system along the same direction — *κ* rescales columns, the direction is fixed. Within-target guide-replicate direction concordance on Jost, compared against a between-target null and a within-guide split-half noise ceiling from the same cell-level bootstraps, tests this directly: strong concordance well above the between-target null and near the noise ceiling is evidence *for* the linear-direction assumption; weak concordance is evidence against it. This diagnostic was not run for this paper and would apply specifically to Jost’s replicate design.

Scope. The perturb-seq-era methods cited in the Introduction that operate in *gene* space (rather than in a low-dimensional program space) — for instance, DAG-, instrumental-variable-, and linear-cyclic-latent approaches restricted to a subset of targeted genes [6–9] — avoid the projection step tested here. The target-held-out failure of the linear-encoding class at d = 30 does not bear directly on those methods; it is a limitation of scope, not a criticism of a different design choice.

### 3.3 Claims versus scope

### 3.4 Limitations

The result concerns three screens, three linear encodings (fixed, footprint, learned) evaluated on the same target-held-out nested-CV recipe, and one estimator family per direction (TSVD / Ridge on the inverse task; Ridge with training-mean baseline on the forward task). Structural alternatives — a diagonal-only, block-modular, symmetric, or GRN-constrained operator; a much smaller d; nonlinear encoders that use covariates beyond target-gene identity; a nonlinear dose-response family; a Perturb-seq design with true replicate targets and calibrated dose titration — were not tested and could behave differently. The identifiability calculation itself, a rank statement about `range(S)`, is correct as a matter of linear algebra and unaffected by the encoding failure.

Both Replogle screens tested here are essential-gene libraries dominated by a shared growth / stress response; whether the result holds for non-essential perturbations is untested. A Replogle genome-wide K562 refit on 200 random non-essential targets would probe this directly and is left to future work.

Two per-screen caveats. Replogle-essential libraries retain one sgRNA per target after the released aggregation step, so within-target replicate diagnostics do not apply to them. Jost has 25 targets, small enough that a target-held-out CV places most of the design outside the training set and the fit sits close to under-determined at d = 30; the d-sweep controls for this at d = 5 but does not turn Jost into a general genome-scale test. Cross-screen quantitative comparison of specific *ρ* values requires care because per-cell noise structure differs across cell lines.

The additive-input model itself is a modelling choice. CRISPRi is closer to a clamp on target transcript than to an additive forcing term. Under a program-space clamp with orthonormal `W` and a rank-deficient `J_gene = W`⋅`J_prog`⋅W^T^, the intervention model is exactly under-identified from projected observations. This is a valid algebraic result under those assumptions rather than a general impossibility; alternative encoder/decoder conventions or explicit gene-space dynamics can change the conclusion. CRISPRi is often promoter-specific, and untargeted alternative promoters can compensate [11], so a knockdown efficiency measured for one transcript may misstate the effective input; the additive-input encoding assumes one input direction per target.

We report every real held-out *ρ* with its across-outer-fold standard deviation and every simulated positive-control *ρ* with its cross-replicate standard deviation. Interpretation carries the same reservation: held-out *ρ* on a real Perturb-seq screen is the quantity that a matched-SNR linear-truth positive control makes calibrated.

## 4. Methods

### 4.1 Framework

For expression `E` ∈ `ℝ ^(n`×`G)` and a control-derived program basis `W` ∈ `ℝ ^(G`×`d)` with orthonormal columns, program coordinates are z = W^T^e. For a guide targeting *g* at efficiency *κ*`_g` ∈ `(0, 1]`, the perturbation input is `u_g = −`*κ*_g W^T^*δ*`_g`; stacking *m* guides gives `U` ∈ `ℝ ^(d`×`m)`. Under the additive-input, steady-state assumption `S = −J`^−1^`U`, so `J`⋅`S = −U`. Right-multiplying by the pseudoinverse identifies the operator’s action on `X = range(S)`:

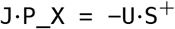

`J` is not identified outside `X`. anchor-op returns this action and blocks access to the full `J` unless both the actuated input subspace `range(U)` and the retained response subspace `range(S)` have rank `d` at the preregistered tolerance (§4.2).

### 4.2 Regularization and the identifiability gate

`S`^+^ is computed by truncated SVD, with the full regularization path retained. A singular direction counts as identified only if σ`_i > rank_tol` ⋅ σ`_max(S)`. The default `rank_tol = 1`×`10`^−2^ was preregistered to prevent eps-based numerical rank from silently accepting below-noise directions on collinear guide libraries. The identifiability gate now labels a fit as full-domain-identified only when both `in-put_rank(U)` and the effective retained response rank equal `d` at the same tolerance; noise-inflated retained rank on rank-deficient `U` was previously accepted and is not. Regression tests cover both the noise-inflated failure mode and the heavy-Tikhonov collapse.

### 4.3 Efficiency estimation

Three estimators with an auto-router. `mean_ratio` (*κ* = 1 − mean_pert / mean_ctrl) is asymptotically unbiased under Poisson and consistent under independent zero-inflation. `poisson_mle` matches under pure Poisson. `detection_rate` (Pr[X_ctrl > 0] − Pr[X_pert > 0]) is not a *κ* estimator on count data but is a valid signed distributional-shift statistic on pre-scaled residual data. The `“auto”` router chooses `detection_rate` when ≥ 2% of entries are negative (the pre-scaled-residual signature), else `mean_ratio`. On Replogle-essential’s pre-scaled residual matrix the router chooses `detection_rate`; on Jost 2020’s UMI counts it chooses `mean_ratio`. Where the manuscript reports a per-screen *κ* distribution, Replogle values are the proxy score and Jost values are the count-based estimate; they are not commensurable on a common fractional-knockdown axis and are not compared as such.

### 4.4 Per-entry noise anchors

Within-guide cell-level split-half bootstraps on each screen: for each target with ≥ 20 cells, split cells into equal halves, compute half-Δz vectors d_1_, d_2_ in that screen’s own PCA basis fit on non-targeting controls, and take the median per-entry standard deviation 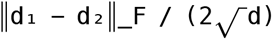 across targets. This gives σ = 0.24 (K562), 0.35 (RPE1), and 0.066 (Jost, per sgRNA). The three anchors are used in every matched-SNR positive control (§4.6).

### 4.5 Target-held-out nested cross-validation

Outer folds: 5 target-grouped folds. All sgRNAs of a target are held out together, so no sibling sgRNA leaks between train and test. Inner folds within each outer training set: 3 target-grouped folds by the same rule. The TSVD rank is chosen per outer fold from the grid `{0, 1, 2, 3, 5, 8, 12, 16, 20, 25, 30}` by minimizing pooled inner-MSE of `A`⋅`S_val + U_val`. `r = 0` returns `A ≡ 0` and gives outer-fold *ρ* = 1 exactly, which lets nested CV select “shrink to zero” whenever the fitted model does worse than predict-zero on inner validation. We refit at the selected rank on the full outer training set and report pooled *ρ* across outer folds with the across-outer-fold standard deviation.

The Jost d-sweep in §2.3 truncates S and U to the first `d’` ∈ `{5, 10, 15, 20, 25, 30}` rows before running the same recipe. We ran the d-sweep on Jost only: the two Replogle-essential screens train on ~150 targets at d = 30 and are already comfortably overdetermined, so a sweep in d would not add information there. Jost with 25 targets is the design in which under-determination at d = 30 is a live concern, and the sweep is what rules it out. The K562 random-panel distribution in §2.3 rebuilds the entire pipeline once (variance-based highly-variable-gene selection with target genes force-kept in the feature matrix, PCA on non-targeting controls at d = 30, per-guide response construction with the proxy-efficiency path used in the released essential-gene notebooks) on the 1,581 K562 essential targets that pass the pipeline’s retention filter (≥ 60 cells plus per-target guide-response QC in `measure_operator`), saves Σ`_ctrl` and `W` from that single fit, and then subsamples 50 random 200-target panels from the saved measurement to compute panel-level nested-CV *ρ* under the fixed Σ`_ctrl` and basis. 4.5b Comparator panel: fixed, footprint, and learned linear encodings The comparator panel of §2.6 evaluates three encodings on the forward task under target-grouped nested-CV (5 outer folds × 3 inner folds, same as §4.5). *Fixed*: Ŝ_ `test = B`⋅`U_test + b` with `U_g` = W^T^*δ*`_g`, `B` fit on training with ridge (λ chosen from a log grid `{0, 10`^−3^, …, 10^6^, 10^30^} on in-ner folds; λ = 10^30^ collapses `B` → `0` and yields the training-mean intercept). *Footprint*: same as fixed with U_g = W^T^ Σ`_ctrl` *δ*`_g`. *Learned*: a training-fold learned linear embedding following the one-table convention of Ahlmann-Eltze et al. [10] with a training-fold-mean intercept. Within each outer training set we compute gene-space pseudobulk responses for each training target, SVD them to `d_embed = 30`, and use `V_gene[:, gene_index[target]]` — the target-gene column of the training-derived gene loading matrix — as the perturbation embedding for both training and test targets. We verified this implementation against a Python port of the published linear-baseline routine. An earlier two-table implementation deviated by 7 % on the K562 pseudobulk fidelity test and was replaced before the analyses reported here; the code, test, and superseded outputs are preserved in the versioned repository. Ridge is fit on the centered training responses; prediction adds the training-mean intercept back in gene space and projects through `W` for the program-space metric (the gene-space metric is also reported). Baselines: predict-zero (*Ŝ*= 0, *ρ*_fwd = 1) and predict-training-mean (column mean of training `S` broadcast). Nulls for the learned encoding: shuffled-embedding null (≥ 100 permutations of the target→embedding map, refit per permutation, intercept preserved). Preregistered success (per screen, per encoding): *ρ*`_fwd` more than 2 outer-fold SDs below the training-mean baseline AND (learned only) below the 2.5th percentile of the shuffled-embedding null.

### 4.5 c Positive-control ensembles at matched amplitude

The Supplementary Fig S2 ensembles of §2.2 rerun the matched-SNR linear-truth control across five ground-truth families. *Dense*: `J_ref = G − 1.5`⋅`I` with `G` entrywise `N(0, 1/d)`. *Sparse-10 / 2%*: a Bernoulli mask at 10% / 2% is applied off-diagonal to `G` before the `−1.5`⋅`I` diagonal. *Rank-5*: `J_int= B`⋅`V` with `B` ∈ `ℝ^(d`×`5)`, `V` ∈ `ℝ^(5`×`d)` both entrywise Gaussian scaled by 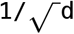, plus the same diagonal. *Block-modular*: 5 blocks along the diagonal, per-entry Gaussian at 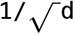 within blocks and 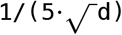) between blocks. α`_S` is re-derived per (screen × ensemble) so the median column norm of `S_true` matches the observed median. Draws with spectral abscissa of `J_true` above zero are rejected and re-drawn; `N` ≥ `15` accepted draws per cell.

#### 4.6 Matched-SNR linear-truth positive control

For each screen we compute the observed median column norm of `S`. For each of 15 independent simulations, we draw `J_ref = G − 1.5`⋅`I` with `G` entrywise `N(0, 1/d)` and set `J = J_ref /` α`_S`, choosing α`_S` so the median column norm of the noise-free `S_true = −J`^−1^`U` matches the observed median at each screen’s own `U`. We add per-entry noise at the screen’s bootstrapped σ (§4.4) and run the same nested-CV recipe (§4.5) on this synthetic `(S_obs, U)` pair. We report the mean and standard deviation of held-out *ρ* across the 15 simulations.

The interaction-only cosine reported in §2.2 uses the same synthetic ground-truth ensemble at matched α_S and reports Frobenius cosine after subtracting the diagonal from both `A` and `J`, with a cross-replicate null as its baseline. The cross-replicate null pairs each fitted `A_r` with an independently drawn ground truth `J_{r’}` (shift-1 pairing in the replicate index; N = 200 replicates), so every element of the pipeline is preserved and only the ground truth we score against is randomized. The signal-scale sweep in §2.2 varies α_S across a log grid and reports the cos-versus-α_S curve and the α_S implied by the observed median column norm.

### 4.7 Direction-only refit

Column-normalize `S` and `U` — divide each column by its own Frobenius norm — then run the same nested-CV recipe (§4.5). This removes all per-guide scale from the fitting problem. For the matched-SNR linear-truth control we generate `S_obs = S_true +` σ ⋅ ε at each screen’s own α`_S` and σ on the same U, column-normalize both `S_obs` and `U` in the same way, and run the same recipe. Both are reported in §2.3.

### 4.8 Preregistered vs post hoc

Preregistered: `rank_tol = 1`×`10`^−2^, TSVD regularization, `d = 30` PCA basis fit on non-targeting controls, the raw bin-split `rel_diff` ≤ `0.25` threshold. Post hoc analyses declared in `PREREGIS-TRATION_AMENDMENT.md` before running: per-screen σ anchors (§4.4, replacing an earlier common σ); the α_S signal-scale sweep and matched-α positive control (§4.6); the target-grouped nested-CV recipe and the rank-0 baseline (§4.5); the interaction-only cosine analysis (§4.6); the direction-only refit (§4.7); the K562 random-panel refit and the Jost d-sweep (§2.3, §4.5); the identifiability gate now requiring `rank(U) = d` as well as retained `rank(S) = d`, a bug fix affecting the identifiability label on rank-deficient inputs with no numerical-recovery changes; the footprint-encoding preregistration of 2026-09-27; the Steps 1–3 preregistration of 2026-09-28 covering the comparator panel of §2.6 and §4.5b, the positive-control ensembles of §4.5c, and the K562 random-panel distribution of §2.3; and the learned-encoding fidelity-check preregistration of 2026-09-28 that triggered the one-table convention fix in §4.5b. All amendments are shipped with the release; results are reported as they came out.

### 4.9 Data availability

Replogle et al. 2022: gwps.wi.mit.edu / Figshare Plus deposit 20029387 (K562 essential sampled at day 6; RPE1 essential sampled at day 7) [1]. Jost et al. 2020: GEO GSE132080 [2].

### 4.10 Code availability

Source code is available under the MIT License at the anchor-op repository, release v0.3.2 https://github.com/manarai/Anchor-op. A pinned conda `environment.yml` (Python 3.11), a 59-test pytest suite, per-figure regeneration scripts, and the learned-encoding fidelity script (`reproduction/54_learned_encoding_fidelity.py`, a Python port of `solve_y_axb` from Ahlmann-Eltze et al. 2025’s linear-baseline code) are shipped with the release, together with every JSON that a numeric claim traces to. Every numeric claim in this manuscript that is not derived in the text has an entry in `results/recheck/number_provenance.csv` giving the value stored in a JSON in `results/recheck/`, the reproduction script that produced it, and a note on rounding where applicable. Round-form numbers cited in the abstract and Results are drawn from the un-rounded values in that table.

## Author contributions

Kyler Fullmer: Validation, Visualization, Writing — review & editing.

Dalton Kutzen: Methodology, Formal analysis, Investigation, Writing — original draft, Writing — review & editing.

Tommy W. Terooatea: Conceptualization, Supervision, Project administration, Funding acquisition, Writing — review & editing.

## Funding

This work was supported by the College of Life Sciences at Brigham Young University.

## Competing interests

The authors declare no competing interests.

## AI use disclosure

The authors drove the main scientific idea and the coding analysis. Generative AI was used to assist analysis and to edit the manuscript text (drafting, revising, and consistency passes). All numeric results reported here were produced by the reproduction scripts in `reproduction/` and traced in `results/recheck/number_provenance.csv`; the authors verified the analyses and take responsibility for the final content of the manuscript.

## Supplementary Figures

**Supplementary Figure S1.**
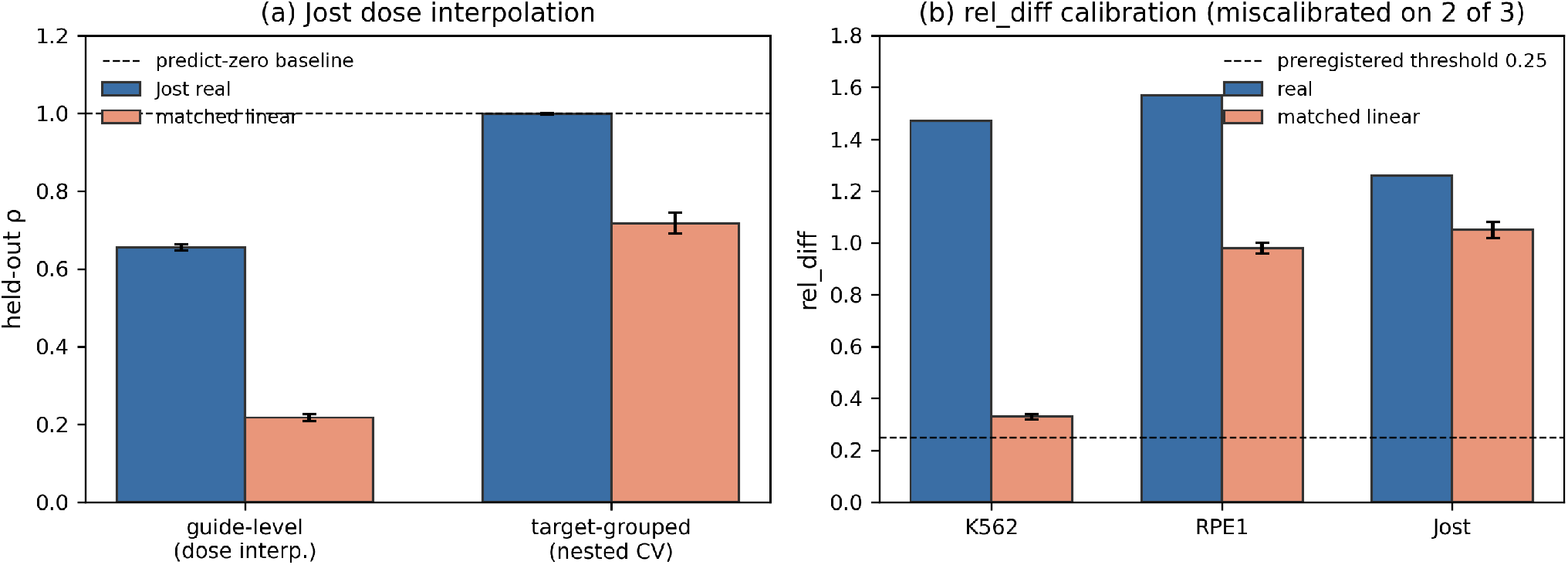
*(a)* Jost dose interpolation: guide-level held-out *ρ* (siblings in training) is 0.66 for real Jost vs 0.22 for matched linear-truth (SD 0.008); under the same target-grouped nested crossvalidation used in Table 1, real Jost sits at the predict-zero baseline (*ρ* = 1.00, 5-fold SD 0.002) while the matched linear truth reaches *ρ* = 0.72 (SD 0.027, N = 15). The gap between the two settings shows the guide-level *ρ* is dose interpolation along a known target direction rather than operator-level recovery on a new perturbation target. *(b)* `rel_diff` against a matched linear ground truth. The preregistered0.25 threshold is unreachable even for a linear truth on RPE1 and Jost, so it is miscalibrated at these signal-to-noise ratios; compared with its matched control, real `rel_diff` is higher on all three screens (K562 1.47 vs 0.33; RPE1 1.57 vs 0.98; Jost 1.26 vs 1.05), consistent with the held-out failure.

**Supplementary Figure S2.**
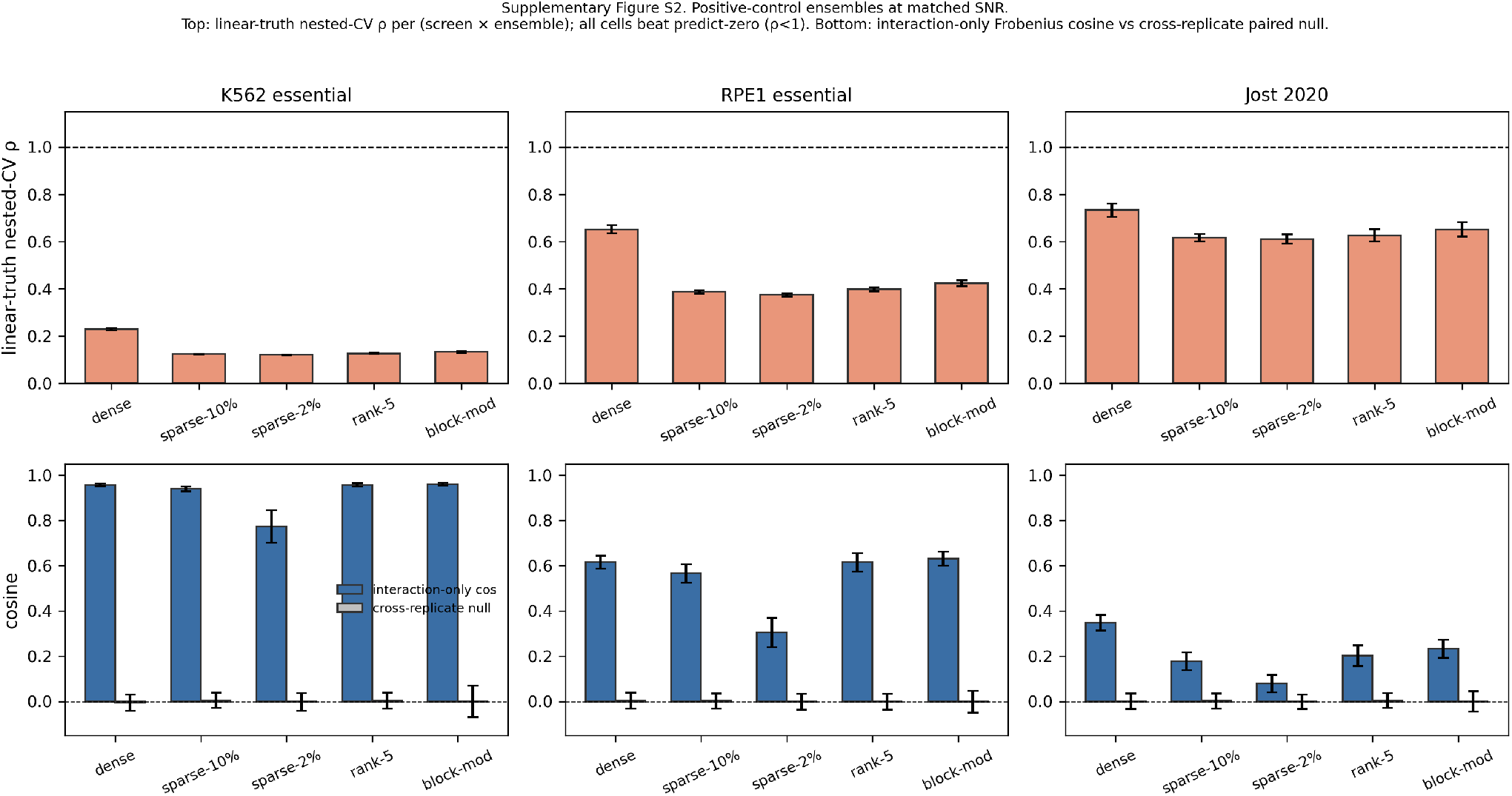
Positive-control ensembles at matched SNR. *(top row)* Linear-truth nested-CV *ρ* per (screen × ensemble); all cells beat predict-zero (*ρ* < 1) across dense, sparse-10 %, sparse-2%, rank-5, and block-modular ground truths on K562 essential, RPE1 essential, and Jost 2020. *(bottom row)* Interaction-only Frobenius cosine vs the cross-replicate paired null.

## Notes

https://plus.figshare.com/articles/dataset/_Mapping_information-rich_genotype-phenotype_landscapes_with_genome-scale_Perturb-seq_Replogle_et_al_2022_processed_Perturb-seq_datasets/20029387

https://www.ncbi.nlm.nih.gov/geo/query/acc.cgi?acc=GSE132080

### Competing Interest Statement

The authors have declared no competing interest.

